# High-throughput development and characterization of new functional nanobodies for gene regulation and epigenetic control in human cells

**DOI:** 10.1101/2024.11.01.621523

**Authors:** Jun Wan, Abby R. Thurm, Sage J. Allen, Connor H. Ludwig, Aayan N. Patel, Lacramioara Bintu

## Abstract

Controlling gene expression and chromatin state via the recruitment of transcriptional effector proteins to specific genetic loci has advanced the potential of mammalian synthetic biology, but is still hindered by the challenge of delivering large chromatin regulators. Here, we develop a new method for generating small nanobodies against human chromatin regulators that can repress or activate gene expression. We start with a large and diverse nanobody library and perform enrichment against chromatin regulatory complexes using yeast display, followed by high-throughput pooled selection for transcriptional control when recruited to a reporter in human cells. This workflow allows us to efficiently select tens of functional nanobodies that can act as transcriptional repressors or activators in human cells.

## Main text

The synthetic manipulation of gene expression and epigenetic state has become a widely useful technique for studying transcriptional regulatory networks and developing molecular tools to reprogram cells or combat disease. One way to implement this type of control is to fuse transcriptional effector domains or epigenome editors to DNA-binding domains that target specific genomic loci^1–4^. This approach allows us to better understand transcription and chromatin, dissect the connection between phenotype and genotype, and create gene regulatory networks. However, in many cases, entire chromatin regulatory complexes must be recruited to a genomic locus in order to achieve a maximal effect^5,6^. These complexes and their individual members are quite large, and therefore not ideal candidates for synthetic experiments or disease therapies due to limitations on packaging in viral vectors and cell delivery. Moreover, overexpression of chromatin regulators via direct delivery often causes cytotoxicity^7^, emphasizing the potential power of indirect recruitment of chromatin regulatory complexes for genetic manipulation.

We have shown before that an effective alternative strategy for transcriptional and epigenetic control is to recruit nanobodies against endogenous chromatin regulators (CRs) to genomic loci of interest^8^. Nanobodies are small, single-domain antibodies known for their tight binding affinities and versatile uses in biology that range from live-cell imaging to next-generation anticancer therapeutics^9–11^. Structurally, nanobodies resemble the VH domain of classical antibodies, consisting of constant regions interspersed by three hypervariable complementarity-determining regions (CDRs) that determine each nanobody’s specificity in target recognition and binding affinity^12^. They are produced naturally by camelids (llamas, alpacas, and relatives) and are therefore normally isolated from animals injected with a protein cocktail that contains the antigen of interest^13^. This method is time- and resource-intensive and severely limits the throughput by which nanobodies against different targets can be produced. Recently, a yeast display platform for expression and selection of a large nanobody library presented on the surface of yeast has been developed and made widely available, massively increasing the amount of nanobodies now screenable for use^14^. This library has already been used successfully to identify nanobodies that bind therapeutically relevant proteins such as GPCRs, the SARS-CoV-2 Spike protein, and immune checkpoint inhibitor CTLA4^15,16^.

Despite these recent advances in nanobody production and selection, there exist very few nanobodies that are known to bind chromatin regulators^8^. Importantly, it is also difficult to predict which member of a large CR complex must be bound in order to most effectively capture and control the entire complex. Moreover, even when good binders are found via yeast display, they need to function inside mammalian cells. Naive nanobody libraries (consisting of ∼10^8^ members^14^) are intractable for direct testing in human cells due to technical limitations that arise from culturing enough cells to have sufficient representation of each library member; to this end, it is necessary to curate a smaller starting library of nanobodies more likely to affect mammalian gene expression for testing in cells.

In order to circumvent this bottleneck and screen directly for nanobodies that can have an effect on gene expression, we combined yeast display of a synthetic nanobody library^14^ against chromatin regulatory complexes that can act as corepressors or coactivators with HT-Recruit, a pooled reporter recruitment assay that tests for transcriptional repression or activation in mammalian cells^1^ (**Fig. 1A**). This workflow begins by overexpressing a FLAG-tagged chromatin regulator in HEK-293 cells, immunopurifying the chromatin regulator complex from cell lysates using a FLAG antibody, and directly using the purified complex as bait in the yeast display selection assay. (**Fig. 1A, top, Methods)**. The naive synthetic nanobody library (consisting of at least ∼10^8^ unique and diverse members^14^) is grown up in yeast, with each yeast cell expressing and displaying a specific nanobody on the cell surface, and subjected first to one round of negative selection against unconjugated magnetic beads to remove non-specific binders. The remaining yeast-displayed nanobodies are subjected to several rounds of positive selection against magnetic beads that have the FLAG-tagged chromatin regulator bound to their surface. These positively selected nanobodies are then amplified and cloned as fusions to the doxycycline (dox)-inducible reverse tetracycline (rTetR) DNA-binding domain (**Methods**). This resulting library (reduced in size closer to 10^4^) is then used in a pooled high-throughput transcriptional recruitment assay^1^, in which each of the rTetR-nanobody fusions is recruited to a constitutively transcribing (active/ON) or non-transcribing (silenced/OFF) reporter gene downstream of TetO binding sites. The reporter gene encodes both the fluorescent protein Citrine and a surface reporter that can be used for magnetic separation of large cell populations, allowing for facile separation of ON and OFF populations and classification of nanobodies as potent transcriptional activators or repressors (**Fig. 1A, bottom**).

**Figure 1.**
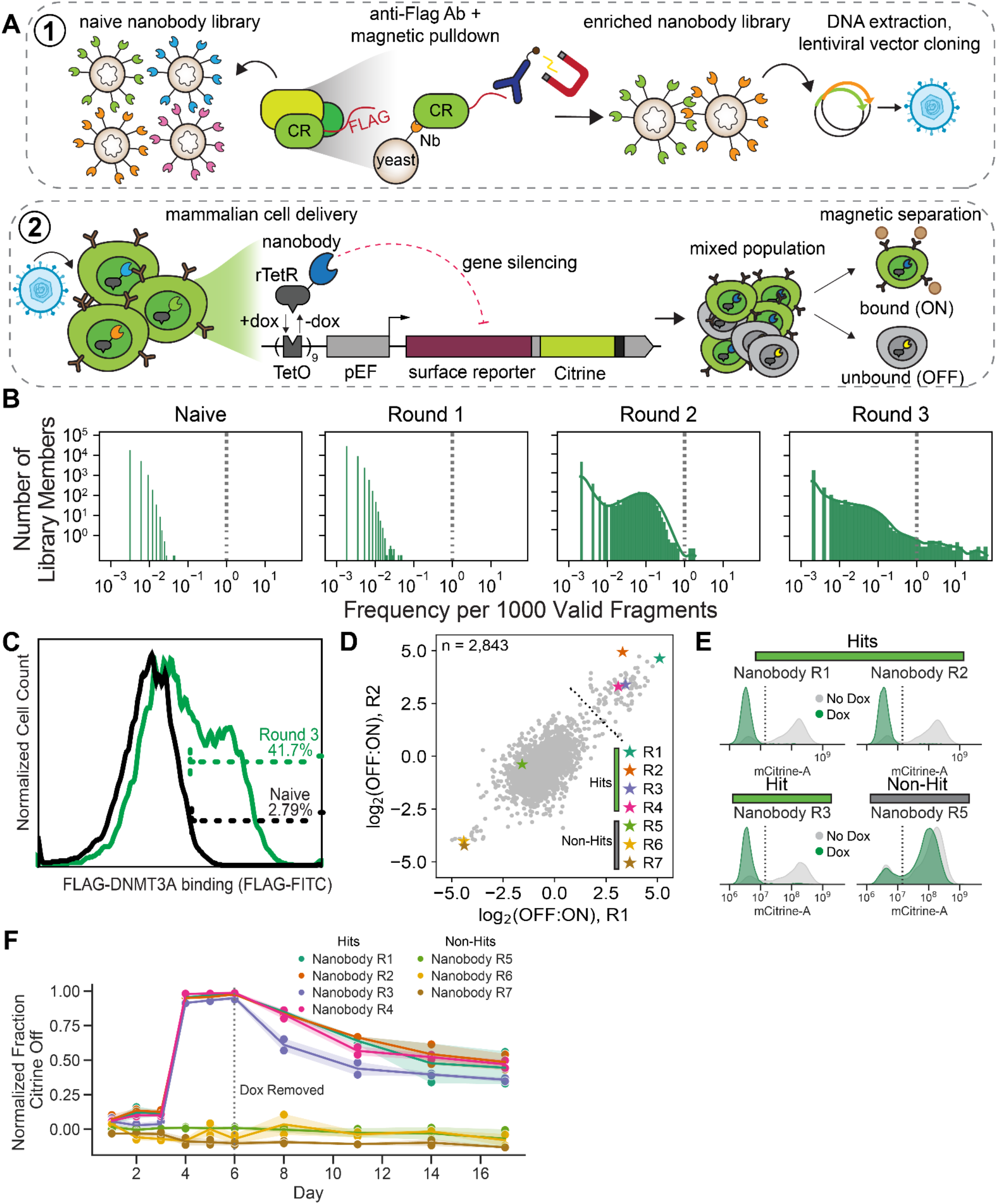
A yeast display nanobody library contains members that silence a reporter gene in human cells. (A) Schematic of library enrichment from yeast display and assay in human cells. Top, yeast display: a large naive nanobody library is genetically encoded in yeast, and each yeast cell expresses a single nanobody displayed on its surface. FLAG-tagged chromatin regulator complexes (CR) are overexpressed in human cells, which are then lysed and mixed with nanobody-displaying yeast.The library is enriched through multiple rounds of magnetic pulldown, and the final library is sequenced. DNA is extracted from yeast and directly cloned as fusions to a doxycycline (dox)-inducible rTetR DNA-binding domain. Bottom, HT-recruit: next, human cells encoding a reporter gene with a Tet-responsive element, fluorescent marker, and surface expression marker are infected at low MOI with rTetR-nanobody lentivirus. Cells are binned into silenced (reporter gene off, surface marker not expressed) and non-silenced (reporter gene on) pools by magnetic separation against the surface marker, and nanobody library member frequencies in each pool are computed. (B) Distributions of library member frequencies in the naive yeast library and after each round of selection. X-axis, frequency per 1000 valid sequenced library members, y-axis, count, vertical dashed line, frequency of 1 per 1000 fragments. (C) Flow cytometry distribution of yeast cells bound to FLAG-tagged DNMT3A (stained with FITC-conjugated anti-FLAG antibody) in round 3 (green line) compared to naive (black line). (D) Log_2_(OFF:ON) (positive = more silenced) enrichment scores plotted for each replicate of the high-throughput nanobody recruitment screen (R1 and R2) in HEK-293A cells on day 5 of dox-mediated recruitment. Stars indicate nanobodies selected for validation. Hits are defined using an enrichment score threshold of 2. (E) Representative flow distributions for selected nanobodies on day 6 of dox recruitment. Grey curves, no dox added; green curves, dox added. (F) Timecourse of dox-mediated gene silencing and epigenetic memory of selected nanobodies, plotted as the normalized fraction of silenced cells (Citrine Off).

For selecting repressive nanobodies, we chose to perform the yeast display selection against DNA methyltransferase 3A (DNMT3A) based on its known ability to permanently silence genes, both endogenously and upon synthetic recruitment, by depositing methylation at CpG sites on DNA^17–20^. Recruitment of DNMT3A to induce DNA methylation has been shown to increase the potency of epigenetic silencing tools^21,22^, but direct overexpression of DNMT3A is often toxic in mammalian systems^23^. To circumvent this cytotoxicity and develop nanobodies that can recruit endogenous DNMT3A complexes, we first expressed DNMT3A fused to a FLAG-tag in HEK-293 cells, lysed these cells and immunopurified protein complexes using protein G magnetic beads conjugated with anti-FLAG antibody (**Methods 1.1&2**). These magnetic beads with DNMT3A protein complex bound to their surface were mixed with the yeast-display library during three rounds of positive selection. Before each round of positive selection against DNMT3A-coated beads, yeast cells were mixed with beads coated with the FLAG-tag only to remove yeast-expressing nanobodies that bound nonspecifically to the magnetic beads or the FLAG tag. After three rounds of positive selection against DNMT3A-FLAG, the yeast library showed significant enrichment for certain nanobody sequences: many members of the final round 3 library are represented at much higher frequencies than in the naive library or earlier rounds (**Fig. 1B, Table S1**). After round 3 of selection, more than 40% of the yeast cells stained positive when incubated with cell lysate containing DNMT3A-FLAG and fluorescently labeled anti-FLAG antibody (Methods 1.5), implying this library contains a higher numbers of nanobodies able to bind DNMT3A compared to the naive yeast library which contained <3% positive cells (**Fig. 1C**).

The nanobody library enriched against DNMT3A-containing complexes (round 3) was cloned as a fusion to rTetR and delivered via lentivirus to a HEK-293A cell line containing the dual surface marker-Citrine reporter (**Fig 1A, bottom**). Nanobodies were recruited to the TetO sites in the reporter gene by addition of dox for 5 days, after which magnetic separation was performed to isolate OFF cells (containing nanobodies that silenced reporter expression) from ON cells. We extracted genomic DNA from these cell populations and used next-generation sequencing of the nanobody sequences to calculate enrichment scores in the OFF population compared to the ON population. 2,843 nanobody sequences passed the threshold for sequencing depth in both biological replicates (**Methods**). The screen was reproducible between replicates and showed a wide range of enrichment scores (**Fig. 1D, Table S2**). We chose a hit threshold of 2 based on the distribution of scores showing a large cloud centered around zero (likely non-hits) and a tail at high OFF:ON scores (likely hits). To verify this hit threshold, seven nanobodies were chosen for individual validation via flow cytometry, including four hit nanobodies with high log_2_(OFF:ON) enrichment screen scores and significant potential to act as strong repressors, and three non-hit nanobodies: one with a medium to low enrichment score, and two with low enrichment scores (**Fig. 1D, colored stars**). The seven nanobodies were individually cloned into the same rTetR-fusion lentiviral vector and delivered individually to HEK-293A cells, and dox was added for six days to recruit the nanobodies to the reporter gene. The flow-cytometry measurements with these nanobodies matched the screen results showing 100% of cells silenced at day 6 for the four top screen hits (**Fig. 1E**) and good correlation between screen scores and the fraction of cells with Citrine OFF **Fig. S1A**). Overall, we identified 87 nanobodies with screen scores above 2 that act as transcriptional repressors in human cells. Notably, <3% of the yeast-display enriched nanobodies (87 nanobodies out of 2,843 sequences enriched through yeast display) were effective repressors in human cells. That means that if we had randomly selected nanobodies after the yeast display and individually tested them in human cells, it would have been difficult to identify even a handful of successful repressors without testing hundreds of cell lines, demonstrating the utility of high-throughput mammalian cell recruitment assays coupled to yeast display enrichment to efficiently and systematically identify functional nanobodies from a large pool of binders.

We determined the consensus sequence for each variable CDR for all hit nanobodies (those with scores > 2 in the high-throughput recruitment screen) and aligned the top 4 hits to this consensus using the MEME suite motif finder. We found that the validated hit nanobodies (R1-R4) align moderately well to each other and to the consensus for each CDR, while non-hit validated nanobodies R5-R7 show less conservation to the consensus sequence, particularly in CDR3 (**Fig. S1B, Methods**).

Finally, after six days of dox-mediated recruitment of the individual nanobodies to the Citrine reporter, we removed dox to test the nanobodies for effects on epigenetic memory: their ability to induce silencing that is maintained after their release from the target gene (**Fig. 1F**). The fraction of cells with Citrine OFF (silenced cells) was computed for up to 17 days, and the four top screen hits all led to retention of epigenetic silencing in at least 40% of the cells. These results show that at least some of the nanobodies selected against the epigenetic regulator DNMT3A confer some degree of epigenetic memory.

We next wanted to test if our yeast display and mammalian cell screening platform could also be used to identify nanobodies that bind to and recruit coactivators. We chose TET1/2/3 as baits in the assay due to their known ability to directly increase transcriptional activity at an epigenetically silenced locus through oxidative demethylation of 5mC ^5,6,24–26^. TET1 and its cognate DNA dioxygenases TET2 and TET3 have been implicated in the regulation of cell fate determination in development, myelin repair pathways, and NF-kB pathway modulation in cancer, demonstrating their promiscuous ability to affect overall gene expression^27–29^. Additionally, nanobodies that activate epigenetically silenced promoters could be useful in correcting aberrantly methylated and therefore silenced genes, such as in imprinting disorders in development or tumor suppressor gene silencing in cancer^11^.

To establish a system that could screen for functional nanobodies that reactivate an epigenetically silenced gene, we first established a silenced reporter by transiently recruiting an rTetR-DNMT3A fusion to our reporter. We transfected rTetR-DNMT3A into HEK-293A cells harboring our TetO site-surface marker-Citrine reporter, and added dox at the time of transfection to recruit the DNMT3A before it was diluted out of the cells (**Fig. 2A**). We then sorted a silenced (Citrine OFF) population that remained stably silenced in the absence of any recruitment (**Fig. 2B**), and could be reactivated by recruitment of the catalytic domain of TET1 (TET1-CD), which has previously been used as an effective and stable de-methylator for activation of silenced genes^26^ (**Fig. S2A**).

**Figure 2.**
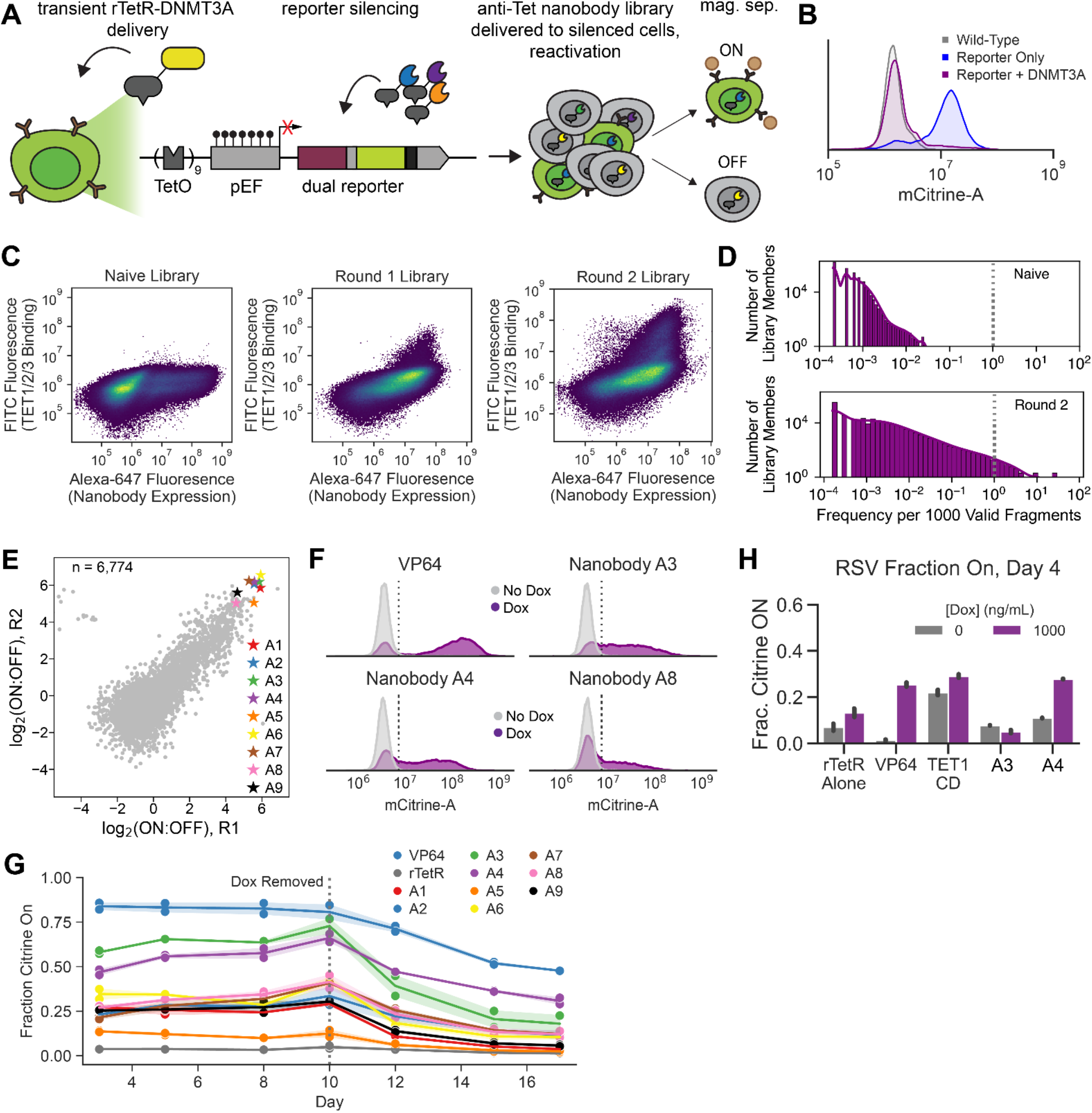
Yeast display selects nanobodies that activate silenced genes. (A) Schematic of silenced reporter and activation screen. Human DNMT3A was fused to rTetR and transfected transiently into HEK-293A cells harboring the TRE-surface marker-citrine reporter while dox was added. Silenced (citrine OFF) cells were sorted and a stable population was maintained. Yeast display was performed using full length human TET1, mouse TET2 and human TET3 to curate a library of nanobodies against TET1/2/3 complexes, which were fused to rTetR, delivered via lentivirus to the silenced cells, and separated into re-activated (bound) and still-silenced (unbound) pools. (B) Flow cytometry measurements of Citrine expression wild-type HEK-293A cells (gray), HEK-293A cells expressing the surface marker-Citrine reporter (blue), and HEK-293A cells expressing the reporter after transient transfection and recruitment of rTetR-DNMT3A (purple). The silenced population (purple curve) was sorted and maintained for screening use. (C) Flow cytometry measurements of the naive library (post-negative selection), round 1-TET enriched, and round 2-TET enriched nanobody-expressing yeast libraries. X-axis, Alexa-647 fluorescence (indicating nanobody expression on the yeast surface); y-axis, Alexa-488 fluorescence from anti-FLAG antibody (indicating yeast binding to FLAG-tagged TETs). (D) Distributions of library member frequencies in the naive library (top) and round 2 yeast display selection against the TETs. X-axis, frequency per 1000 valid sequenced library members; y-axis, count; vertical dashed line, frequency of 1 per 1000 fragments. (E) Log_2_(ON:OFF) (positive = more activated) scores per replicate of the TET nanobody recruitment screen in HEK-293 cells; nanobodies chosen for follow-up denoted as stars. (F) Flow cytometry Citrine measurements for selected nanobodies and VP64 on day 10 of dox-mediated recruitment. VP64 was chosen as an activating positive control. Grey curves = no dox added, purple curves = dox added. (G) Timecourse of dox-mediated reporter activation starting from day 3 of dox addition, plotted as the fraction of cells that have re-activated (citrine ON). Lines show the average of two biological replicates, dots are individual replicates, and shading shows standard deviation. (H) Measurements of the fraction of cells activated (Citrine ON) upon recruitment of rTetR alone (negative control), VP64 (positive control), TET1-CD (the catalytic domain of TET1 enzyme fused to rTetR), and activating nanobodies #A3 and #A4 to epigenetically silenced promoter RSV. Grey, no dox; purple, dox added. Bars indicate the average of two biological replicates, and error bars show standard deviation.

To produce an enriched anti-TET nanobody library, we performed 2 rounds of yeast display enrichment with anti-FLAG-antibody beads coated with FLAG-TET1, FLAG-TET2 and FLAG-TET3 (in combination) (**Methods**). Good enrichment of both nanobody surface expression and FLAG-TET binding were seen after each round of positive selection compared to both the naive library (post-negative selection) and glucose-exposed control (**Fig. 2C-D, Fig. S2B, Table S3**).

This anti-TET-enriched nanobody library was cloned and delivered to the silenced reporter cells. Doxycycline was added for 5 days to recruit the anti-TET nanobodies to the reporter, and cells were subsequently sorted into ON (reactivated) and OFF (silenced) populations using magnetic separation as before (**Fig. 2A**). The activation screen in silenced HEK-293A cells was reproducible for the 6,774 nanobodies that passed the sequencing depth threshold (**Fig. 2E, Table S4**). Nine nanobodies that were among the highest enrichment scores in the screen were chosen for follow-up (**Fig. 2E, colored stars**), and again individually cloned into the rTetR-fusion vector before being delivered to HEK-293A cells with the stably silenced reporter (described above). For individual validations, we used rTetR fused to VP64 as a positive control, which consists of 4 repeats of the potent trans-activating domain VP16 from herpes simplex virus 1 (**Fig. 2F**). The empty rTetR-fusion vector was used as a negative control (**Fig. S2C**). Most of the nanobodies chosen show some activation by day 10 of dox recruitment, with two nanobodies (A3 and A4) showing strong activating potential close to that of VP64 (**Fig. 2F-G**). The rTetR only vector did not activate any of the silenced cells, proving that the activation by VP64 and the screened nanobodies is not simply due to a DNA binding domain facilitating reactivation by binding near the promoter (**Fig. S2C**).

To investigate the dynamics of gene re-activation and test for any epigenetic memory of activation that occurred upon recruitment of the nine selected nanobodies, we continued to measure Citrine expression for 10 days of dox treatment and 7 days after dox removal (**Fig. 2G)**. By two days after dox release, the percentage of cells active had already decreased; the most strongly activating nanobodies also decreased quickly but maintained some significant fraction (>0.25) of cells activated at day 17. VP64, the most potent activator, maintained the highest level of activation memory at day 17, which is surprising since it is not known to recruit DNA demethylases directly, but instead has been shown to activate by the strong recruitment of general transcription factors (eg. TFIID) and chromatin remodelers (e.g. BAF)^19^. The loss of activation after release of the anti-TET nanobodies suggests that nanobody recruitment is not sufficient to permanently remove all epigenetic marks associated with the silencing of the reporter gene.

We also tested two of the top-performing nanobodies (A3 and A4) in K562 cells, where we recruited to TetO sites upstream of RSV, a viral promoter that is rapidly silenced in both HEK-293A and K562 cells, and minCMV, a minimal promoter. Neither nanobody could activate the minCMV promoter above background (**Fig. S2D**), while nanobody A4 (but not A3) could activate the RSV promoter to levels similar to VP64 (**Fig. 2H**). This implies that the activation activity may be specific to epigenetically silenced (not minimal) promoters, but may also have some dependence on the mode of epigenetic silencing of the target promoter (DNMT3A recruitment vs. endogenous silencing of a viral promoter).

Although the mechanism by which these nanobodies act warrants further investigation, it is clear that the yeast display coupled to the mammalian high-throughput testing was successfully able to identify nanobodies that can act as repressors or activate epigenetically silenced promoters, and that are more compact than endogenous chromatin regulator complexes. These tools will be useful both for studying mechanisms of background silencing in synthetic contexts and for activating silenced genes that affect cell phenotype genome-wide. The combination of large-scale yeast display and mammalian high-throughput screening, techniques that have been developed and refined for use only in recent years, has the potential to massively accelerate the rate at which new protein tools and therapies are discovered and understood. Our system in particular, which leverages a diverse synthetic nanobody library with recruitment of protein complexes to genetic loci, offers a chance to expand the capabilities of live-cell imaging, synthetic gene circuit design, and gene therapy through the ability to recruit important protein complexes easily to a desired gene. Expanding this platform to target endogenous genes or coupling it with fluorescent reporters for other processes and their matching protein baits will allow development of functional nanobodies for many biological applications.

## Supporting information

Table S3. CDR sequences and counts for each round of anti-TET1/2/3 nanobody enrichment.

Table S1. CDR sequences and counts for each round of anti-DNMT3A nanobody enrichment.

Table S4. CDR sequences and population counts for high-throughput anti-TET nanobody recruitment activation screen in human cells

Table S2. CDR sequences and population counts for high-throughput anti-DMNT3A nanobody recruitment repression screen in human cells

## Author Contributions

J.W. and L.B. conceptualized the study. J.W. performed yeast display and yeast immunostaining experiments. J.W. performed high-throughput screens in mammalian cells with assistance from C.H.L., and performed some of the initial individual flow cytometry validations. A.R.T. and S.J.A. performed the individual flow cytometry experiments for the data included in this manuscript with assistance from A.N.P. C.H.L. performed the initial high-throughput data analysis. A.R.T. performed all other data analysis included in the manuscript and made the final figures. A.R.T., J.W., and L.B. wrote the manuscript with input from all others. L.B. supervised the study.

## Acknowledgements

We thank Prof. Mike Bassik, Dr. Josh Tycko, Dr. Mike Van, and Dr. Haotian Du for helpful advice and experimental assistance. This work was supported by NIH-NHGRI R01HG011866 (L.B.), a Stanford School of Medicine Dean’s Postdoctoral Fellowship (J.W.), NIH-NCI F30CA287739-01 (A.R.T.), a Stanford Bio-X Bowes Fellowship (A.R.T.), a Stanford Sarafan Chem-H Chemistry-Biology Interface Fellowship (A.R.T.), and NIH T32GM145402 (A.R.T.).

## Competing Interest Statement

J.W. and L.B. have submitted a provisional patent application related to this work. L.B. is a co-founder of Stylus Medicine and a member of its scientific advisory board. All other authors declare they have no known competing interests.

## Supplemental Information

Table S1. CDR sequences and counts for each round of anti-DNMT3A nanobody enrichment.

Table S3. CDR sequences and counts for each round of anti-TET1/2/3 nanobody enrichment.

## Methods

### Resource and Code Availability

All materials are available from the corresponding author upon request.

## 1. Yeast Display

### 1.1 Expression of FLAG-tagged Chromatin Regulators (CRs) in HEK-293 cells

Full length DNMT3A, or TET1, TET2, or TET3 fused to the 3xFLAG were cloned into a pRetro-CMV2-TO-puromycin vector using Gibson assembly. 24 hours prior to transfection, HEK-293 cells were plated in 4 x 10 cm plates in DMEM (Thermo Scientific 10569044) + 10% FBS (Sigma-Aldrich F0926-500ML) supplemented with L-glutamine and Pen/Strep (Fisher Scientific 10378-016). At the time of transfection, cells were 70-80% confluent. The following day (20-24 hours later), cells were transfected with pRetro-CMV2-TO-3xFLAG-CR (DNMT3A, TET1, TET2 or TET3). Before transfection: in each 10cm plate, medium was changed to 20 mL serum-free DMEM (no FBS, no Pen/Strep). For transfection, two sets of 1.5 ml tubes were prepared: one set containing 450 μl 2X HBS (50 mM HEPES, pH 7.05, 10 mM KCl, 12 mM dextrose, 280 mM NaCl, 1.5mM Na_2_PO_4_) and another set containing 25μg plasmid (pRetro-CMV2-TO-3xFLAG-CR) + 65μl 2M CaCl_2_ into 0.1XTE (450 μl total). DNA and CaCl_2_ were pipette-mixed and added dropwise to the 2X HBS. This mixture was incubated at room temp for 1 min. The DNA-calcium-phosphate co-precipitate was added dropwise to the surface of the media containing the cells and the plate was swirled gently to mix. Ten to twelve hours after transfection, the medium was gently aspirated, and 10 ml of pre-warmed DMEM medium (+10% FBS, no Pen/Strep) was added without disturbing the fine precipitates on the bottom of the plate. 48-72 hrs post transfection, transfected cells were harvested.

### 1.2 Preparation of CR-coated magnetic beads for yeast display

HEK-293 cells transfected with each individual CR were scraped in 10 mL lysis buffer (50 mM Tris–HCl, pH 8.0, 1 mM EDTA, 150 mM NaCl, 1% NP40, 1× PMSF, and 1 x NEM) and cell lysates kept on ice for 30 min. Cell lysates were cleared by centrifugation at 10,000 × *g* for 1 min and the supernatant was kept on ice. To prepare magnetic beads for immunoprecipitation, anti-FLAG magnetic beads (Bimake B26102) were resuspended in the vial via pipette-mixing. 150 μL (the amount may be scaled up or down as required) of the bead suspension was transferred to a new tube with 0.5 mL TBS buffer (50 mM Tris HCl, 150 mM NaCl, pH 7.4). The mixture was gently pipetted 5 times. The tube was placed on the magnet for 10 seconds to separate the beads from the solution after which the supernatant was discarded. This step was repeated 2 times. (Note: Prepare all magnetic beads together and then divide into aliquots if processing multiple samples). To bind the CRs complex to the magnetic beads, ∼500 μL of cell lysate was added to the washed magnetic beads. The tubes were gently rotated for 2 h at room temperature or overnight at 4°C. The tubes were placed on the magnet to separate the beads from the solution for 2 minutes and the supernatant was transferred into a new tube. (Note: During the binding process, magnetic beads occasionally cluster together). To wash any non-specifically bound proteins from the beads, 500 μL PBST was added to the tube (136.89 mM NaCl; 2.67 mM KCl; 8.1 mM Na_2_HPO_4_; 1.76 mM KH_2_PO4; 0.5% Tween-20), and the magnetic beads were resuspended by pipetting gently. The tube was rotated for 5 min, and placed on the magnet to separate the beads from solution for 2 minutes to remove the supernatant. This wash step was repeated 2 times. If contaminating proteins are still bound non-specifically to the beads (as measured by Western blot), extend the wash time, increase the number of washes, or increase the detergent content in the wash buffer.

### 1.3 Yeast growth and induction

The yeast nanobody library^14^ was maintained in Yglc4.5 -Trp medium (1 liter: 3.8 g of -Trp drop-out media supplement (US Biological), 6.7 g Yeast Nitrogen Base, 10.4 g. Sodium Citrate, 7.4 g Citric Acid Monohydrate, 10 mL Pen-Strep (10,000 units/mL stock), and 20 g glucose, pH 4.5). Nanobody expression is under the control of the GAL1 promoter: nanobodies are only produced on the cell surface when the yeast is grown in a galactose-containing medium. Expression of the nanobody library was induced by dilution of a yeast aliquot of ≥10x library diversity into -Trp +galactose medium (1 liter: 3.8 g -Trp drop-out media supplement (US Biological), 6.7 g Yeast Nitrogen Base, 10 mL Pen-Strep (10,000 units/mL stock), 20 g glucose or galactose (glucose for normal growth and galactose for induction of nanobodies), pH 6) followed by shaking for 48 hours, at 25 °C, 220 rpm. For the initial yeast dilution, we used at least 5×10^10^ yeast cells in the inoculum to ensure >10x coverage of the nanobody library during each passage and to avoid loss of nanobody clones in passaging.

### 1.4 Bead-Based Enrichment (BBE) of nanobodies using yeast surface display

CR-coated beads were prepared as described in step 1.2 for positive enrichment. 150 μl washed beads were removed from the magnet, resuspended in 1000 μl ice-cold selection buffer (20 mM HEPES, pH 7.5, 150 mM sodium chloride, 2% (w/v) BSA, 1 mM EDTA), and placed on ice until needed. 5×10^10^ induced yeast were used for the first round of selection and 5×10^8^ induced yeast were for subsequent rounds. Yeast were washed and resuspended in selection buffer and then incubated with the CRs antigen-coated beads at 4°C for 2 hours. This yeast display step was conducted against the TET1, TET2, and TET3 proteins together, by combining the coated beads from all three tubes in step 1.2 with induced yeast. Yeast display against DNMT3A was performed separately.

#### 1.4.1 Perform yeast negative selection

Each round of BBE selection began with a negative selection preclear step. Yeast were incubated with non-antigen-coated beads to remove yeast-expressing nanobodies that bound nonspecifically to the magnetic beads. Specifically, 150 μL resuspended magnetic beads conjugated with anti-FLAG-antibody and coated with FLAG tag alone (no CR) were added to the yeast cells induced with galactose. To produce these FLAG-coated beads, we first transfected HEK-293 cells with the pRetro-TO-FLAG empty vector to express the FLAG tag only, and 48 hours post-transfection, we performed immunoprecipitation (IP) using lysates from these cells. We then used these FLAG-coated-beads for negative selection, as follows: Cells were incubated and rotated at 4°C for 2h. Upon completion of the incubation, the tube was placed on the magnet, taking care to transfer any liquid lodged in the cap of the tube to the bottom portion of the tube. After 2 minutes, the supernatant was carefully removed from the tube and transferred into a fresh 10 mL tube labeled “negative preclear #1”. Beads were resuspended in 1 mL ice-cold selection buffer with a pipette and placed on the magnet for 2 minutes. Supernatant was removed and discarded; beads were resuspended in 1 mL ice-cold selection buffer and set aside. The pre-clear step was repeated using the supernatant from the previous step as input, and then the resulting depleted supernatant was carried through to the following step.

#### 1.4.2 Perform yeast positive selection

After the negative selection, nanobodies were enriched over 3 rounds of BBE selection by staining the yeast with CR complex-coated beads. Specifically, the yeast cells after negative selection were mixed with the complex-coated magnetic beads and rotated at 4°C for 2h. Upon completion of the incubation, the tube was placed on the magnet, taking care to transfer any liquid lodged in the cap of the tube to the bottom portion of the tube. The cells and the beads were incubated on the magnet for 2 minutes. The supernatant was carefully removed from the tube and discarded. The beads were resuspended in 10 mL ice-cold selection buffer using a pipette and then placed on the magnet for 2 minutes. The supernatant was removed. The beads should contain a population of yeast cells containing nanobodies enriched for binding to the target, in this case DNMT3A, or TET1/2/3.

#### 1.4.3 Rescue the enriched yeast population

Beads from the previous positive selection were resuspended in -Trp4.5 media and transferred to a sterile culture tube containing 4 mL -Trp4.5 media (5 mL -Trp media, beads, and cells in total). The tube was vortexed gently and 5 μL sample was collected from the culture. The 5 μL sample was diluted into 995 μL -Trp4.5 media (200x dilution) and set aside in a clean tube labelled “positive #1” for a later analysis step. The cells on the beads were grown at 30°C with shaking for 48 hours.

#### 1.4.4 Plate fractions of beads from negative and positive selectionsorts to estimate the number of cells recovered in each step

The saved supernatant from the negative sorts (negative preclear #1) were vortexed, and 100 μL was transferred into 400 μL fresh -Trp4.5_media. The diluted samples were vortexed and 5 μL of each sample was transferred into 995 μL -Trp4.5_media (200x dilution) (tube labeled as negative preclear #2). The 200x dilutions of the negative sort (negative preclear #2) and the positive sorts (positive #1) were vortexed and 10μL from each population was transferred into 190μL -Trp4.5_media (4000x dilution). A -Trp4.5_media plate was divided into four regions using a permanent marker; each dilution was vortexed and 20u was plated. The plate was grown at 30°C for 3 days and resulting colonies counted. One colony in the 200x and 4000x dilutions represents 5×10^4^ and 1×10^6^ cells recovered, respectively. This step allows for an estimate of the library size after each round.

#### 1.4.5 Prepare selected yeast sorted cells for further rounds of selection

After the overnight growth, the cells’ OD600 was measured. If the OD600 is still low, continue to allow the cells to grow; another day of the growth is acceptable in case of especially low OD600s. Once the culture approached saturation, cells were pelleted (at 900xg for 5 minutes) and supernatant aspirated. The pellet was resuspended in 1 mL Trp4.5_media and transferred to a 2 mL tube. The tube was placed on the magnet for 2 minutes. The supernatant was recovered and diluted into two cultures for further expansion. 2.5×10^8^ cells were diluted into 25 mL -Trp4.5_media for growth and induction, and the remaining cells were diluted into 25 mL -Trp4.5_media for overnight growth and temporary storage at 4°C in case the first selected population needs to be induced and selected again. Once the 2.5×10^8^ yeast cell culture containing ∼2.5×10^8^ cells reached an OD600 between 2 and 5, 5×10^8^ cells were pelleted and resuspended in 50 mL -Trp4.5_media. Cells were incubated at 25°C with shaking for 48h to induce.

### 1.5 Confirm CR complex binding to the enriched nanobody library

3xFLAG-tagged DNMT3A or TET1/2/3 was expressed in HEK-293T cells, and the resulting cell lysate containing the DNMT3A complex was used as the selection antigen. After each round of BBE selection, following galactose induction of nanobodies, nanobody-expressing yeast were incubated with the DNMT3A,or TET1/2/3 complex-containing lysate, washed, and then stained with Anti-DYKDDDDK Tag (FLAG tag) Mouse Monoclonal antibody (FITC (Fluorescein)) (GenScript, A01632, 1:50 dilution), and HA-Tag (6E2) Mouse mAb (Alexa Fluor® 647 Conjugate)(Cell Signaling Technology, 3444S, 1:50 dilution). DNMT3A, or TET1/2/3 binding was confirmed and analyzed by flow cytometry (Biorad ZE5) to verify the enrichment for nanobody binders compared to the naive yeast library.

After three rounds of BBE selections, the library of nanobody plasmids was extracted from the enriched yeast library by Zymoprep Yeast Plasmid Miniprep II (Zymo D2004).

## 2. High-throughput screening of nanobodies capable of silencing in human cells

### 2.1 Pooled library cloning of selected nanobodies into a lentiviral construct

The library of enriched yeast nanobody library plasmids was extracted after three rounds of yeast display enrichment and then PCR amplified with Q5 Ultra II Master Mix (NEB M0544L). 10 ng of nanobody library template was added to 23 μl H2O, 1 μl of each 10 μM primer^1^, and 25 μL master mix for 8 50 μL reactions. All primers used for library amplification are available in^1^. Reactions were amplified as follows: 3 minutes at 98°C, then 25 cycles of 98°C for 10s, 55°C for 30s, 72°C for 50s, and a final extension at 72°C for 10 minutes. The resulting amplified dsDNA libraries were loaded onto a 2% TAE gel (run for 25 minutes at 100 V) and excised at the expected length (around 400 bp). A QIAgen gel extraction kit (Qiagen 28706) was used for purification of excised DNA. The libraries were cloned into lentiviral recruitment vectors pWJ036 (anti-DNMT3A nanobody screen) or pWJ254 (anti-TET nanobody screen) with 4x10 μl GoldenGate reactions (75 ng of pre-digested and gel-extracted backbone plasmid, 5 ng of library (2:1 molar ratio of insert:backbone), 0.25 μL of T4 DNA ligase (NEB M0202M), 0.75 μL of Esp3I-HF (NEB), and 1 μL of 10x T4 DNA ligase buffer) using 60 cycles of digestion at 37°C and ligation at 16°C for 5 minutes each, followed by a final 5 minute digestion at 37°C and then 20 minutes of heat inactivation at 70°C.

Golden Gate reactions were pooled together and purified using MinElute columns (Qiagen 28004), eluting the combined products in 6 μL of ddH_2_O. 2 μL of the resulting eluate was transformed into 50 μL of Endura DUO electrocompetent cells (Lucigen 60242-2) and recovered in 2 mL of LB, shaking, for one hour. Cells were then plated on 6 10’’ x 10’’ LB-agar plates with carbenicillin. After overnight growth at 30°C, colonies were scraped off plates and collected. Plasmid pools were extracted with a HiSpeed Plasmid Maxiprep kit (Qiagen 12663). After purification, domains were amplified from the original oligo pool and the plasmid pool using primers with Illumina adapter extensions (as in ^1^) using 10 ng of input pool, 23 μL H2O, 1 of each 10 uM primer, and 25 μL of Q5 Ultra II master mix. Amplification conditions were as follows: 3 minutes at 98°C, then 17 cycles of 98°C for 10 s, 55°C for 30 s, 72 °C for 50 s, and a final step of 72°C for 10 minutes. Sequencing datasets were analyzed as described in section **2.5** to determine the uniformity of coverage and synthesis quality of the libraries. In addition, 20 - 30 colonies from the transformations were Sanger sequenced (Quintara) to estimate the cloning efficiency and the proportion of empty backbone plasmids in the pools.

### 2.2 High-throughput recruitment to measure nanobody silencing activity

For the DNMT3A-enriched nanobody screen, HEK-293T cells were plated on four 10-cm plates to generate sufficient quantities of the lentiviral libraries. 5×10^6^ HEK-293T cells were plated on each plate in 10 mL of DMEM, grown overnight, and then transfected with a mixture of the three third-generation packaging plasmids (6.5 μg pMDLG/pRRE, 5 μg Rev, 3.5 μg VSVG, Addgene #s 12259, 12253, and 12251; all gifts from D. Tronos) and 10 μg of rTetR-nanobody library using the calcium phosphate method. Lentivirus was harvested at 48 hours and 72 hours and filtered through a 0.45-mm PVDF filter (Thermo Scientific 168-0045 ) to remove any cellular debris. 8 10-cm plates of HEK-293 reporter cells (already expressing pJT055, the surface marker-Citrine reporter) were infected with the lentiviral library in two separate biological replicates. Infected cells were grown for 3 days, after the cells were selected with 2 μg/mL puromycin for 3 days. Infection and selection efficiency were monitored every other day using flow cytometry to measure mScarlet-(and thus nanobody) positive cells (using a BioRad ZE5). Cells in each 10-cm plate were transferred to 15-cm plates to increase maintenance coverage to >25,000x cells per library element (a very high coverage level that compensates for losses due to incomplete puro selection, library preparation, and library synthesis errors). On day 3 post-infection, nanobody recruitment at the reporter was induced by treating the cells with 1 μg/ml doxycycline (Fisher Scientific 40-905-0) for 5 days. Cells were split every other day and measured for maintenance coverage using flow cytometry.

A pre-silenced reporter cell line was prepared as follows: two 10-cm plates of HEK-293A cells stably expressing pJT055 (surface marker-Citrine reporter) were transfected using Lipofectamine LTX (following manufacturer’s protocol, Fisher Scientific 15-338-100) with 10 μg of a plasmid expressing rTetR-DNMT3A. Doxycycline was added at a final concentration of 1 μg/ml to the cells concurrently with transfection. 48 hours post-transfection, cells were sorted on a SONY FACS machine to isolate the DNMT3A-silenced (Citrine-negative) population. Doxycycline was removed from the media and Citrine-negative cells were maintained in DMEM as normal to allow residual plasmid to dilute out.

For the TET-enriched nanobody screen, the above lentiviral production was repeated using 2 10-cm plates of anti-TET-enriched nanobody lentiviral libraries to infect 6 10-cm plates of HEK-293A cells silenced transiently with DNMT3A, in biological replicate. Selection and recruitment steps were performed identically to the previous screen.

### 2.3 Magnetic separation of reporter cells

At each timepoint, HEK-293 cells were trypsinized and spun down at 300xg for 5 minutes. Cells were then resuspended in 15 mL of PBS (Thermo Fisher 10010023) and centrifuged again. Protein G Dynabeads (ThermoFisher, 10003D) were prepared by gentle pipetting to resuspend. 50 mL of blocking buffer was prepared per 2 x 10^8^ cells by adding 1 g of biotin-free BSA (Sigma Aldrich A4503) and 200 μL of 0.5 M pH 8.0 EDTA (Thermo Fisher, 15575020) into DPBS (GIBCO), and then kept on ice. 60 μL of beads was prepared for every 1 x 10^7^ cells as follows: beads were added to the magnetic stand, storage buffer was removed, and an equivalent volume of blocking buffer was added. Beads were resuspended via pipetting or gentle vortexing, added back to the magnetic stand where buffer was removed, and finally resuspended in 1 mL of blocking buffer per 200 μL of original bead volume. The prepared beads were used to resuspend the previously prepared cell pellets, and the cell-bead mixture was incubated for 90 minutes on a nutator at room temperature. After incubation, the bead and cell mixture were placed on the magnetic rack for > 2 minutes. The unbound supernatant was transferred to a new tube, placed on the magnet again for > 2 minutes to remove any remaining beads, and then the supernatant was transferred and saved as the unbound fraction. Then, the beads were resuspended in the same volume of blocking buffer, magnetically separated again, the supernatant was discarded, and the tube with the beads was kept as the bound fraction. The bound fraction was resuspended in the original volume of blocking buffer. Flow cytometry (ZE5) was performed using a small portion of each fraction to estimate the number of cells in each fraction (to ensure library coverage was maintained) and to confirm separation based on Citrine reporter levels. Finally, the samples were spun down and the pellets were frozen at -20°C until genomic DNA extraction.

### 2.4 Genomic library preparation and next generation sequencing

Genomic DNA was extracted with the QIAamp Blood Maxi Kit (Qiagen 51192) following the manufacturer’s instructions with up to 1 x 10^8^ cells per column. DNA was eluted in EB and not AE to avoid subsequence PCR inhibition. The domain sequences were amplified by PCR with primers containing Illumina adapters as extensions. A test PCR was performed using 400 ng of genomic DNA in a 50 μL (half size) reaction to verify if the PCR conditions would result in a visible band at the expected size for each sample. Then, 25 x 50 μL reactions were set up on ice. PCR was performed identically to section **2.2**, except 400 ng of genomic DNA was used as input for each reaction. The PCR reactions were pooled and ≥ 140 μL were run on at least three lanes of a 2% TAE gel alongside a 100-bp ladder for at least one hour, the library band around 400 bp was cut out, and DNA was purified using the QIAquick Gel Extraction kit with a 30 μL elution. Libraries were then quantified with a Qubit dsDNA HS Assay Kit (Thermo #Q33231) and Agilent TapeStation (Agilent #G2964AA) and sequenced on an Illumina NextSeq with a 300-cycle High-Output kit using paired-end sequencing (forward read 200 and reverse read 100 cycles) and 8-cycle index reads.

### 2.5 High-throughput sequencing data analysis

Sequencing reads were demultiplexed using bcl2fastq (Illumina). Individual CDR sequences were extracted from read pairs and CDR sequence combination instances were counted using the Python script ‘make_nanobody_counts.py’. Briefly, the script uses portions of the nanobody constant sequences that bookend each CDR to define the boundaries of and extract CDR sequences along with their per-nucleotide quality scores. CDR1 and CDR2 information were extracted from the R1 read while CDR3 information was extracted from the reverse complement of the corresponding R2 read. CDR-wide mean quality scores were computed from the per-nucleotide quality scores, and read sequence and quality information were compiled into a dataframe. Reads with one or more undetected CDR and/or with mean quality scores less than 30 were filtered out. Reads with identical CDR combinations at the DNA-sequence level were grouped and counted. This process was repeated for each sample sequenced. The enrichments for each nanobody (CDR combination) between OFF and ON samples were computed using the script ‘makeRhos.py’. In this script, nanobodies with fewer than 5 reads in both samples for a given replicate were filtered out, whereas nanobodies with fewer than 5 reads in one sample would have those reads adjusted to 5 to avoid inflating enrichment values due to low sequencing depth. Counts were normalized to the sum of counts in that sample to account for differences in sequencing depth (in effect, frequencies were computed) prior to computing log2(OFF:ON) enrichment scores. All code for high-throughput screening analysis can be found on Github at https://github.com/bintulab/HT-recruit-Analyze.

## 3. Individual validations of nanobody function in human cells

### 3.1 Silencing and activation assays measured by flow cytometry

Individual nanobodies were synthesized as gBlocks from IDT and cloned as direct fusions to rTetR using Gibson assembly into the recruitment vectors pWJ036 or pWJ254, as used in the high-throughput recruitment assays. HEK-293 cells expressing the surface marker-Citrine reporter were infected with each nanobody separately. One day after infection, selection for the nanobody constructs was started using puromycin (2 ug/mL), and continued until > 90% of the cells were mCherry positive (∼2 days). Cells were split into separate wells of a 24-well plate and either treated with doxycycline (1 ug/ml) to recruit the rTetR-Nanobody to the reporter or left untreated. After 6-10 days of treatment, doxycycline was removed by spinning down the cells, replacing media with PBS to dilute any remaining doxycycline, and then spinning down the cells again and transferring them to fresh media. Time points were measured every 2-3 days by flow cytometry analysis of > 10,000 cells on a ZE5 flow cytometer (BioRad). Data was analyzed using Cytoflow (https://github.com/cytoflow/cytoflow, courtesy of bpteague). Events were gated for viability and for mCherry as a delivery marker.

### 3.2 Consensus sequence determination for hit nanobodies

The sequences of each CDR for all hit anti-DNMT3A nanobodies (those with enrichment score >2 in the repressor nanobody screen) were submitted to the MEME Suite for consensus construction^30^. Clustal Omega was used for initial multiple sequence alignment (MSA) analyses, and final alignments for selected nanobodies were performed by hand as seen in **Fig. S1A**.

## Supplementary Figures

**Supplementary Figure 1.**
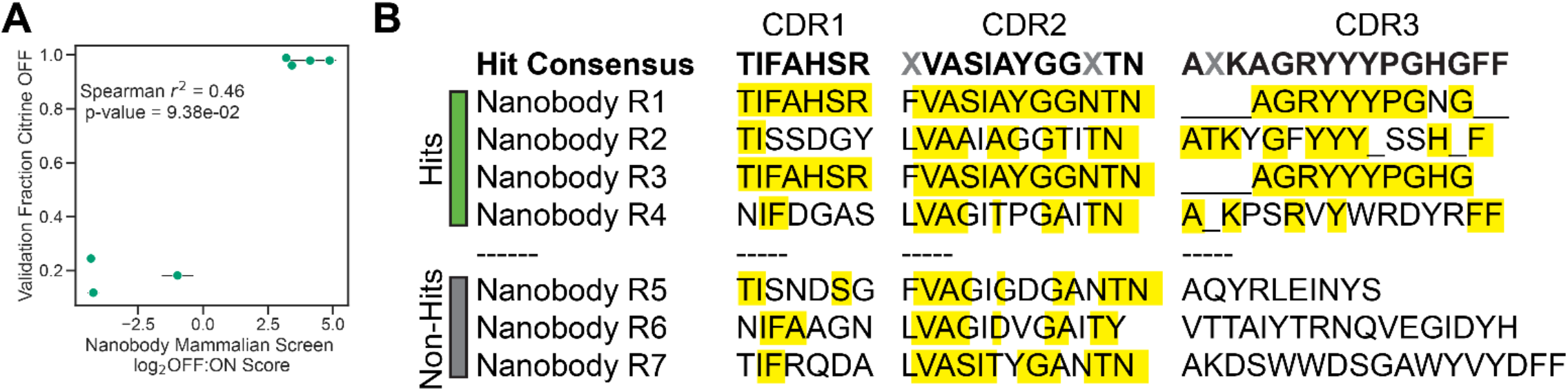
Details of DNMT3A yeast selection and screen, related to Figure 1. (A) Individual flow cytometry measurements of selected nanobodies fused to rTetR, as in **Fig. 1E**, compared to their scores in the high-throughput recruitment assay. X-axis, HT-recruit screen score; y-axis, fraction cells with Citrine OFF as measured by flow cytometry. Error bars indicate standard deviation, and dots represent the average of two biological replicates. (B) Sequences of each of the CDRs (variable regions) of selected hit or non-hit nanobodies, as in **Fig. 1D**, compared to the consensus sequence as computed by the MEME server. Conserved residues in each CDR are highlighted in yellow.

**Supplementary Figure 2.**
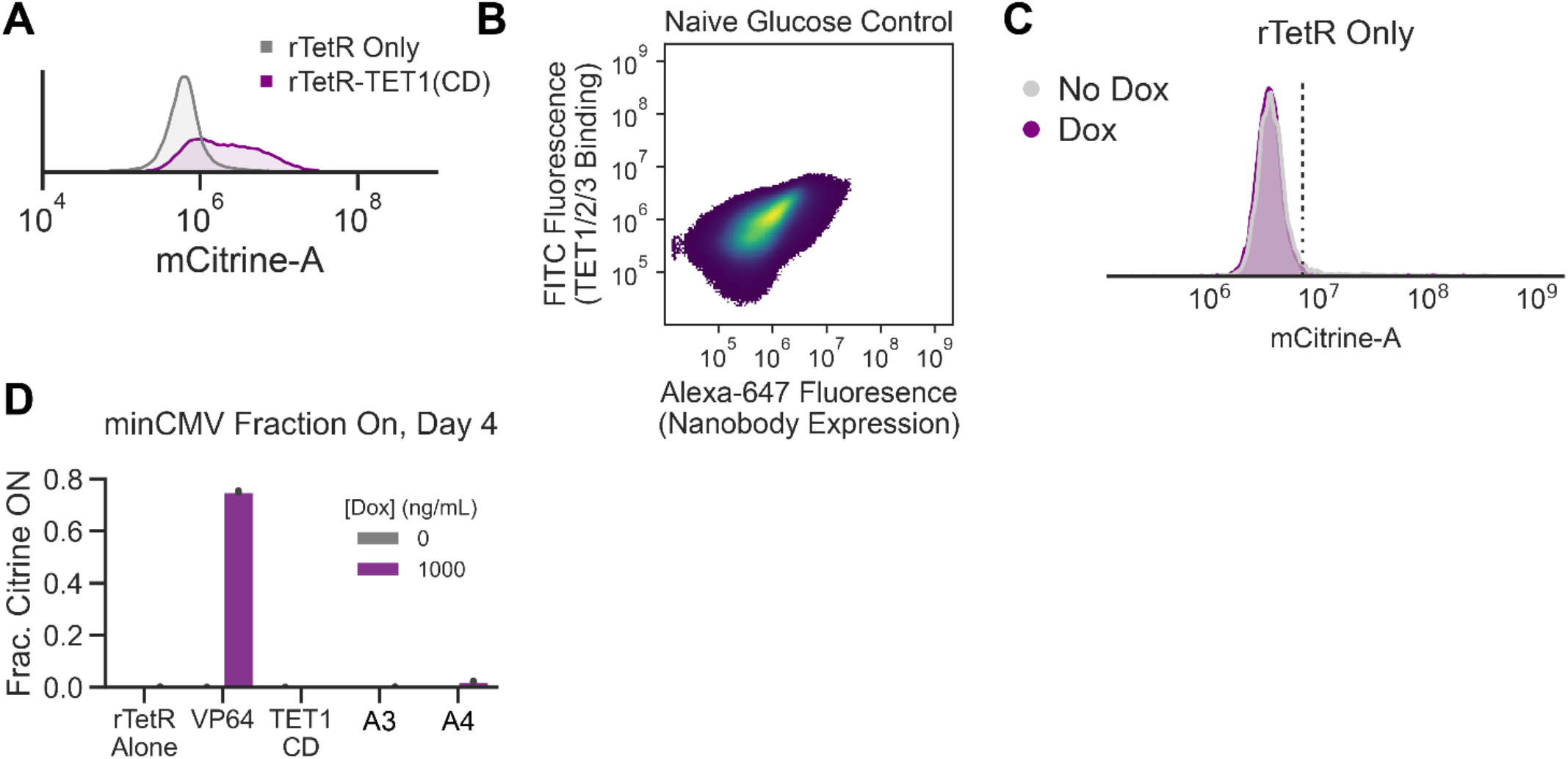
Details of TET yeast display and recruitment screen, related to Figure 2. (A) Flow cytometry measurements of DNMT3A-silenced and sorted HEK-293A cells after 5 days recruitment of rTetR only (gray) or rTetR-TET1(CD). (B) Flow cytometry measurements of a negative control, glucose-exposed yeast library. X-axis, Alexa-647 fluorescence (indicating nanobody expression on yeast surface); y-axis, Alexa-488 fluorescence (indicating yeast binding to FLAG-tagged TETs. (C) Flow cytometry Citrine measurements for rTetR only negative control on day 10 of dox recruitment to the epigenetically silenced pEF reporter. Grey curve = no dox added, purple curves = dox added. (D) Measurements of the fraction of cells activated (Citrine ON) upon recruitment of rTetR alone (negative control), VP64 (positive control), TET1-CD (the catalytic domain of TET1 enzyme fused to rTetR), and activating nanobodies #A3 and #A4 to the minimal promoter minCMV. Grey, no dox; purple, dox added. Bars indicate the average of two biological replicates and error bars show standard deviation.

